# The study of the *Drosophila innubila* Nudivirus in cells and flies

**DOI:** 10.1101/2025.06.06.658368

**Authors:** Margaret E. Schedl, Kent M. Mulkey, Robert L. Unckless

## Abstract

2.

Species of the fruit fly, *Drosophila,* are essential models for investigating host/virus interactions. Research aimed at understanding how hosts mount immune defenses against viruses and how viruses evade those immune defenses relies on host/virus pairs that have evolved together overtime, thus necessitating a need for a DNA virus model system for *Drosophila* with a virus that naturally infects the host. The *Drosophila innubila* Nudivirus (DiNV) is an emerging model system poised to take on the role as the DNA virus model for *Drosophila*. In this paper we describe the development of the DiNV model system with animal and cellular resources. We describe the development of new *Drosophila* cell lines, virus titration assays, anti-nucleocapsid antibodies, and injection-based DiNV infections in *Drosophila innubila*. We find that *D. innubila* cells can be used to grow DiNV for use in further experiments, and that DiNV is highly virulent when injected into adult *D. innubila*. The resources we have developed enable substantial future research on the DiNV-*D. innubila* system.

**Impact statement:** DNA viruses are the cause of several important and challenging human diseases including hepatitis B, mononucleosis, respiratory disease, gastroenteritis, herpes and chickenpox. Recently there has even been a resurgence of many viral infections due to decreasing rates of vaccination, so to develop better therapeutics and intervention strategies, we must better understand the dynamics of how hosts and DNA viruses interact. The development of a natural DNA virus model in *Drosophila* creates the opportunity to better understand host/DNA virus interaction.

**Data summary:** All data and code are available at in our Github repository: available upon publication

## 5. Introduction

Arguably all life on earth is affected by viruses, from microscopic bacteria to the largest plants and vertebrates. In nearly every ecosystem or habitat where scientists look closely enough, viruses are the most abundant biological category (1). In contrast to all other living organisms, there appears to be no universal common ancestor for viruses (1, 2). Viruses affect our food supply, costing approximately $30 billion per year worldwide from crop loss (3). However, there are some instances where viruses and hosts have evolved a mutualistic relationship, notably between some bacteria and bacteriophages (4). Additionally, viral DNA sequences commonly become incorporated into the genomes of macroorganisms. For instance, approximately 8% of the human genome is comprised of endogenous retroviral genetic content (5), and for some species of parasitoid wasps, the presence of viral genomes within the host genome are essential for reproduction (6).

Very early on in the development of model organisms for biological research, the viruses that infect these burgeoning models began to become an important factor. For instance, in the 1920s researchers working with *E. coli* began investigating bacteriophages that infected their bacteria strains (7). In the 1960’s some of the first research on resistance to retroviruses was done on murine leukemia virus in mice (8). Today the use of model organisms for virus research remains extensive and influential. *Drosophila* as a model is well known for its abundance of molecular and genetic tools, which can be readily harnessed for virology research.

Insects like *Drosophila* have only innate immune systems to respond to pathogens and lack the adaptive immune response that is observed in other vertebrates. However, *Drosophila* is still suitable as a model for human pathogens, as the innate immune response in humans and *Drosophila* is remarkably similar (9). Innate immunity is a first line of defense against any pathogen in both humans and flies, while an adaptive response takes many days to generate an effect (10), and using *Drosophila* for research allows for the careful study of just the innate immune response. The more we understand about how viruses combat innate immune systems, the more we can design therapeutics against them. Many of the innate immune pathways in *Drosophila*, like the NF-κB signaling for recognition and protection against bacteria and fungi, dSTING for recognition of DNA in the cytoplasm, and RNA interference mechanisms have direct homologs in humans (11–13), leading to a rich literature on innate immunity in *Drosophila*.

While the *Drosophila* system has been a key component in research on viruses, major questions in the realm of the innate immune response remain unanswered. For instance, if the “innate response [is] the evolutionary memory of the species’’ to a pathogen (9, 14), then some aspects of a *Drosophila* response to a human-infecting virus may be missing their evolutionary context. Indeed, the host/virus arms race can be seen as a major driver of co-evolution during the history of an organism, with over 10% of all protein coding genes in eukaryotes being involved in defense mechanisms (1). In *Drosophila*, the rate of adaptive evolution in immune genes is estimated to be twice the rate of the rest of the coding sequence found throughout the genome (15). Subsequently, *Drosophila* has emerged as a leading model for studying these host/pathogen interactions and evolution (16, 17), and a substantial amount of research has been done on *Drosophila’s* native viruses, most notably *Drosophila* C Virus, Sigma Virus, and Nora Virus (18–23). Using *Drosophila*-native viruses provides the evolutionary context that is necessary for research on why and how the defense and counter-defense mechanisms that hosts and viruses deploy, evolve and operate. Even within *Drosophila* species certain viruses can evolve host-species-specific viral suppressors of immunity (24), thus research on the evolutionary arms race between hosts and pathogens benefits when the host/virus pairs share an evolutionary history. Despite the enhanced biological relevance of these viruses natively infecting *Drosophila*, research in this realm has focused almost exclusively on RNA viruses. Only recently were DNA viruses native to *Drosophila* hosts even discovered (25–28), leaving a gap in the field with no well-established model DNA virus system established in *Drosophila*.

We aimed to use the *Drosophila innubila* Nudivirus (DiNV) to fill that gap by developing resources for studying DNA viruses in *Drosophila*. DiNV is a Nudivirus with an envelope and a helical nucleocapsid. It has a double-stranded, circular DNA genome that is 155,555 base-pairs and includes 107 open reading frames (ORFs). DiNV was initially discovered in infected *Drosophila innubila* but also has been associated with other *Drosophila* species in the wild, including members of the *Drosophila* quinaria group: *D. falleni, D. munda, D. tenebrosa,* and also the more distantly related *Drosophila azteca* (28, 29). Nudiviruses are a group of arthropod-infecting viruses closely related to Baculoviruses, and so far, studied nudiviruses share 32 core genes, 21 of which are also shared with Baculoviruses. Nudiviruses have been found to infect and replicate in the gut, fat body, and reproductive tissues of various hosts, and can be transmitted fecal-orally and sexually (30). Although the route of infection and primary infected tissues have not been determined for DiNV, some evidence suggests fecal-oral infection due to the presence of virions in fecal matter from *D. innubila* (28).

Previous work revealed interesting aspects of DiNV evolution, such as recent selection pressures on viral proteins, including those encoding essential proteins for genome replication like *Helicase* (31). Additionally, the recurrent evolution of two DiNV haplotypes that correspond to high and low virulence has been documented within different wild populations of *D. innubila* (29). These different viral haplotypes elicit different gene expression responses in *D. innubila* hosts during infection, notably in Toll-induced immune genes, suggesting that naturally occurring variation in DiNV strains may exert differing selective pressure on hosts (29). Research on a closely related virus, Kallithea Virus, has uncovered that a viral gene; *gp83*, present in both viruses, inhibits the Toll pathway and induces the IMD pathway in hosts during infection (32). These are two main immune pathways in *Drosophila* that canonically produce antimicrobial peptides and continue to be implicated in host response to viral infection even though the mechanism of induction or antiviral effects are not known (18, 32, 33). While some influential research has already been done investigating the dynamic interactions between DiNV and *D. innubila*, much of this research has been done on wild caught infected flies or with DiNV from wild sources where experiments lack some of the necessary controls to draw clear conclusions about DiNV infection.

Here we describe the development of new *Drosophila* cell lines, virus titration assays, anti-nucleocapsid antibodies, and injection-based DiNV infections in *Drosophila innubila*. These laboratory-based resources will allow for more detailed and experimentally based research questions to be carried out on this system. The goal of this resource-building effort is to develop DiNV and *D. innubila* into a dsDNA virus model system, currently lacking in the *Drosophila* field.

## 6. Methods

### Fly lines utilized and fly husbandry

The *D. melanogaster MyD88* mutant line (KG03447) was maintained on molasses food and kept in a 12-hour light, 12-hour dark cycle in a constant temperature room at 22°C. The *D. innubila* line was kept on instant *Drosophila* media (Carolina 4-24 Instant Drosophila Medium, Plain, Item #: 173200, Carolina Biological Supply Co, NC, USA) supplemented with a ∼1g piece of white button mushroom (*Agaricus bisporus)* and a dental cotton roll for pupation. *D. innubila* were maintained in the same 12/12-hour cycle at 20°C.

### Development of embryonic cell lines

Our initial attempts to grow and passage DiNV in available cell lines were unsuccessful (Mulkey *et al.* in prep), so we worked to develop two new cell lines: one derived from *D. melanogaster* carrying an immune gene mutation, and another cell line from wildtype *D. innubila* - the focal host of DiNV. Generation of primary cells from *Drosophila* usually involves crushing developing embryos into clumps or single cells, plating them to a culture flask and waiting to see if any of the primary cells generated survive and grow over long periods of time. The first cell line we developed is from a *D. melanogaster* line with a mutation in the *Myd88* gene that results in a nonfunctional Myd88 protein (genotype: y[1] w[67c23]; P(y[+mDint2] w[BR.E.BR]=SUPor-P)Myd88[KG03447]). This fly line was generated in 2002 by the Berkeley Drosophila Genome Project and is available at the Bloomington Drosophila stock center (Indiana, USA), stock ID 14091. The second cell line was from DiNV’s main host species: *Drosophila innubila*. The fly line chosen was the *D. innubila* genome line because the complete sequenced *D. innubila* genome is available (34). While DiNV does appear to infect other species of *Drosophila*, *D. innubila* is infected in the wild at high frequency (28), and so it seemed like an ideal species to develop a cell line for DiNV growth.

KG03447 *D. melanogaster* flies approximately 2-4 days post emergence were placed in a single 100mm mating cage with an apple juice agar plate bottom. The agar plate was intermittently dolloped with a yeast paste. The collection cage was set up in the morning, and approximately 27 hours after adding flies to the cage, the agar plate was removed. Embryos were removed from the agar plate by rinsing the plate with 1X embryo wash and brushing the plate with a paint brush. The wash was filtered through a 100 μm mesh strainer to retain only the embryos. In contrast, *D. innubila* much prefer to mate in a small 35 mm mating cage instead of the 100mm mating cage, and we used only *D. innubila* aged to 8-11 days post emergence for generating embryos in the mating cages. An average of 50 male and female *D. innubila* genome line flies were added to each mini embryo collection cages (59-105 Embryo Collection Cage-Mini, Genesee Scientific Co., NC, USA) with an apple juice agar plate at the bottom. Eight replicate cages were set up for each collection. Because *D. innubila* push their embryos into the agar, each embryo was removed from the agar with a toothpick under the microscope, and the embryos from each plate were combined in a beaker of 1X embryo wash. The wash was filtered through a 100μm mesh strainer to retain only the embryos.

The subsequent process to generate primary cells from the embryos was the same for both species. The embryos were rinsed through the mesh and resuspended in a conical tube with 10 mL of 50% bleach/1X embryo wash. The embryos were then bleached for 10 minutes to sterilize them and remove the chorion. After the 10 minutes, the solution was centrifuged for three minutes at 400 rpm to pellet the embryos. The following procedure was done in a sterile cell culture safety hood with sterile technique. The embryo pellet was washed three times with serum free Schneider’s *Drosophila* medium containing 100 units/mL penicillin, 100 units/mL streptomycin, 10 μg/mL gentamicin, and 250 ng/mL amphotericin B. Each wash consisted of pipette removal of the supernatant, resuspension of the embryos in 10mL of the wash, followed by centrifugation for three minutes at 400 rpm. Following the third wash, the embryo pellet was resuspended in 2mL of Schneider’s *Drosophila* medium with 10% Fetal Bovine Serum, and antibiotics (same concentrations as above). The solution was then transferred to a sterile 2mL glass Dounce homogenizer and embryos were allowed to settle before homogenizing approximately ten times with moderate force. Homogenization breaks up the embryos into single cells and cell clumps. The homogenized embryos were transferred to a Corning 25cm^2^ plug seal cell culture flask, and the volume increased to 10mL with Schneider’s *Drosophila* medium with Fetal Bovine Serum (10%) and penicillin, streptomycin, gentamicin, and amphotericin B. The harvested cell slurry was split to 2 flasks, 5mL each. The flasks containing the primary cells were incubated at 23°C and allowed to settle and grow. We repeated this process several times to get one flask that produced cells that proliferated continuously and eventually immortalized. This process eventually produced a *D. innubila* cell line, named Dinn-1, and a KG03447 *D. melanogaster* cell line named E-Myd88[KG03447].

### Transfection of newly developed Cell Lines

We aimed to test whether our newly developed cell lines, E-Myd88[KG03447] and Dinn-1, were able to be transfected with a green fluorescent protein (GFP) reporter plasmid. There is not an available plasmid with a GFP gene controlled by a constitutive promoter from *D. innubila*, so pAc5.1B-EGFP (Addgene #21181, MA, USA) was chosen because it contains the gene encoding GFP, with a *D. melanogaster* Actin 5C promoter, and ampicillin resistance. *E. coli* harboring the pAc5.1B-EGFP plasmid were grown in an 150mL overnight culture, and plasmids were extracted using the Qiagen Plasmid Midiprep Plus kit (Qiagen #12941) using manufacturer’s specifications. The plasmid DNA was then quantified with a Qubit Broad Range DNA assay following the manufacturer’s protocol (Thermo Fisher Scientific #Q32850, MA, USA). For transfection, the plasmid DNA was concentrated to 1μg/μL using Ampure XP Beads (Beckman Coulter Life Sciences #A63881, IN, USA). Plasmid DNA was stored at -20°C until use.

The set up for the transfection experiment included three different cell lines. First, we used Dv-1 cells (DGRC Stock 40 (35)), derived from *D. virilis* first instar larvae, as a control for an established cell line that we knew already could be transfected. We also used our two new cell lines, E-Myd88[KG03447] and Dinn-1 cells, to test as well. Dinn-1 cells were plated in 6 wells of a 12 well plate and were seeded at a high density to ensure confluence in the wells as these cells cannot be counted with a hemocytometer. E-Myd88[KG03447] cells were counted with a hemocytometer and were seeded at a density of approximately 200,000 cells per well in three wells of the 12 well plate for 24 hours before transfection. Dv-1 were also counted with a hemocytometer and were seeded at a density of approximately 100,000 cells per well in three wells of the 12 well plate for 24 hours before transfection. Transfection was carried out with the lipofection-based reagent TransIT-2020 Transfection Reagent (Mirus Bio #5404, WI, USA) following manufacturer’s suggested protocols.

### Inoculation of cells in culture with DiNV

We aimed to test whether the E-Myd88[KG03447] cells and the Dinn-1 cells could be infected with DiNV by inoculating cells with the virus and comparing the number of viral genomes produced after five days of incubation. For these experiments an established cell line, Dv-1, was used as a control. Also, for both experiments we used a passage level 4 DiNV isolate produced in a persistently infected Dinn-1 cell line. This virus stock was titered to about 10^5.38^ FFU/mL, and we refer to it as DiNV Passage 4 or P4 (adapted virus, Mulkey *et al.* in prep).

E-Myd88[KG03447] cells were seeded in a 24-well plate at a density of approximately 200,000 cells per well and the Dv-1 cells were seeded at a density of approximately 100,000 cells per well. The cells were plated at different densities because the E-Myd88[KG03447] cells grow slower than the Dv-1 cells. For each cell line there were four replicate wells as a cell control for day 0 and day 5 time points, and there were four replicate wells that received DiNV Passage 4 for both the day 0 and the day 5 timepoint. The cells were allowed to settle to the well bottom for 24 hours before inoculation. The control wells received 10 μL of cell culture medium. The DiNV wells received 10 μL of P4 DiNV. The plate containing the day 0 samples was frozen at -80°C immediately after inoculation. The other plate was placed in a 23°C incubator for 5 days, and then frozen at -80°C.

For the experiment testing the DiNV growth in Dinn-1 and Dv-1 cells in a 24 well plate, only the Dv-1 cells were counted in a hemocytometer. The clumpiness of the Dinn-1 cells does not allow for them to be counted, so they were seeded at a high density to ensure confluence in the wells. Again, the Dv-1 cells were plated at a density of 100,000 cells per well. The cells were allowed to settle to the well bottom for 24 hours before inoculation. The control wells received 10 μL of cell culture medium. The DiNV wells received 10 μL of DiNV Passage 4. The plate containing the day 0 samples was frozen at -80°C immediately after inoculation. The other plate was placed in a 23°C incubator for 5 days, and then frozen at -80°C.

Cells thawed from being frozen at -80°C release from the surface of the cell culture plate and partially lyse. The frozen plates were thawed on ice and 50 μL mixtures of cell and supernatant solution were aliquoted into 1.5mL tubes for each well. Each sample was spiked with 1 μL of a 1:1000 dilution of Lambda DNA to use as a DNA extraction control in qPCR since cell count will vary by sample and cell type. DNA was extracted from each sample using the Qiagen Puregene kit in a modified protocol. In brief, 250 μL of chilled cell lysis solution was added to each sample on ice and pipette mixed and then incubated at 65°C for 10 minutes. Samples were cooled to room temperature and 2μg of RNase A solution was added to each tube. Samples were mixed by inverting 25 times and incubated for 30 minutes at 37°C. After incubation, 100 μL of protein precipitation solution was added to each sample, and samples were vortexed and incubated on ice for five minutes. Then samples were centrifuged for three minutes at 16,300xg to pellet proteins. The supernatant was transferred to new 1.5mL tubes containing 300 μL of fresh 100% isopropanol. Samples were inverted to mix 50 times and centrifuged at 16,300xg for five minutes to pellet the DNA. The DNA pellet was washed once with 70% ethanol, dried, and resuspended in 20 μL of DNA hydration solution. DNA was allowed to resuspend for a few hours at room temperature before being stored at -20°C while waiting for further processes.

The ability of these cells to be infected and replicate the viral genome was tested by estimating the number of viral genomes in each sample using qPCR. Three replicates of each sample were run for qPCR with different primer sets. The E-Myd88[KG03447] experiment samples were amplified with the DiNV *PIF3* gene primers, and primers for the Lambda DNA. The *D. innubila* host gene primer *tpi* was used to quantify the Dinn-1 cell relative genome copy number and the host gene primer *Rpl11* was used for the Dv-1 cell samples. Lambda DNA was spiked into samples prior to DNA extraction and used as an extraction control. This also allowed us to use Lambda DNA Cq values as a proxy for volume. See Table S1 for primer sequences (28). Each reaction included 1 μL of template DNA, 5 μL of SsoAdvanced Universal SYBR Green Supermix (BioRad 1725270), 0.5 μL of 10 μM forward primer, 0.5 μL of 10 μM reverse primer, and 3 μL of molecular grade water. The qPCR program used included: An initial denaturation step of 3 minutes at 95°C, followed by 40 cycles of 10 second at 95°C and 30 seconds at 60°C, followed by a 5-minute 95°C extension. This is then followed by a melt curve section that increases by 0.05°C every 5 seconds from 55°C to 95°C.

Analysis of qPCR results was performed in R (36). We used the formula: ΔCq = (ave Lambda or housekeeping gene Cq - ave PIF3 Cq) and the relative viral genome per volume or cell = 2^ΔCq^. We calculated the genome increase using the 2^ΔΔCq^ method (37) where ΔΔCq = (avg lambda or HK.120hr - avg PIF3.120hr) - (avg lambda or HK.0hr - avg PIF3.0hr) and the relative increase in viral genome from day 0 to day 5 = 2^ΔΔCq^. “HK” refers to the housekeeping gene (*tpi* for *D. innubila* and *Rpl11* for *D. virilis*).

Relative viral genome increase (2^ΔΔCq^) was visualized with ggplot2 (38) with box plots. Statistical differences between relative genome increase were calculated with linear models in R Studio, and we compared the Dv-1 and E-Myd88[KG03447] cell experiment separately from the Dv-1 and Dinn-1 cell experiment. In each model we tested the effect on the non-transformed ΔΔCq and used the R package AICcmodavg to compare models. We compared an ∼infection alone model, an ∼infection + cell type model, and an ∼infection + cell type + infection by cell type interaction models for both datasets. The model with the interaction had the best predictive power for the Dv-1 and E-Myd88[KG03447] cell experiment and was used for analysis. This model was also the best for the Dv-1 and Dinn-1 cell experiment normalized to host genomes. However, the ∼infection alone model was the best fit for the Dv-1 and Dinn-1 cells normalized to Lambda DNA.

### Antibody development against the viral capsid protein (vp39)

To aid in the study of DiNV, we developed a monospecific rabbit polyclonal antibody against the viral nucleocapsid protein, vp39 (DiNV_CH01M_ORF92). This work was performed by Pacific Immunology (Ramona, CA). A peptide sequence (DGRREIISNSNDQTFRPIHRPVIC) matching a viable antigenic target on the vp39 protein was synthesized. A New Zealand White Rabbit was immunized four times with the peptide emulsified with Freund’s complete or incomplete adjuvant. Serum from the initial bleeds was tested by ELISA to confirm immunization and an aliquot of serum from the final bleed was collected and a portion was affinity purified.

The antibody was tested for specificity to DiNV vp39 by Western blot. Proteins from a DiNV Passage 5 stock grown in Dinn-1 cells were separated by polyacrylamide electrophoresis (10%) under reducing conditions as described by Laemmli (39) and transferred to an Immobilon membrane (Millipore) by western blotting using a Bio Rad Protean mini-cell. A lane of Chameleon 700 Pre-stained Protean Ladder (LI-COR, NE, USA) was included on each gel. After transfer the membrane was blocked with DPBS, 2% BSA and 0.1% Tween-20 for 1 hour at room temperature. The blocking solution was removed, and the membrane was soaked in anti-vp39 antibody diluted 1:2000 and mouse anti-Actin (JLA20-s, Developmental Studies Hybridoma Bank, IA, USA) diluted 1:500 in DPBS, 2% BSA and 0.1% Tween-20 and incubated for 1 hour at room temperature. Then, the membrane was washed 3x with DPBS followed by a 1-hour incubation in DPBS, 2% BSA and 0.1% Tween-20 containing goat anti-rabbit IRDye 800CW (LI-COR, NE, USA) diluted 1:5000 and goat anti-mouse IRDye 680RD (LI-COR, NE, USA) diluted 1:5000. After incubation with the conjugates the membrane was washed 3x with DPBS and visualized with a Licor Odyssey M imaging system. The image was processed with FIJI using the Quick Figures plugin (40).

### Fluorescent focus-forming unit (FFU) assay to quantify infectious virus

We aimed to develop a titration method for the quantification of DiNV containing fluids. Note that the standard plaque assay relies on having a virus that causes cytopathic effect in cell culture. Unfortunately, we have not seen that DiNV causes visible cytopathic effect in any cell line that we have used. In contrast, the fluorescent focus-forming unit assay uses only antibody-based detection of virus and is well suited for viruses that do not cause visible effects on cells (41). A fluorescence focus-forming unit (FFU) titer is calculated with this equation:

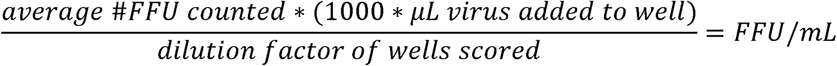

The DiNV FFU method was validated by titration of two new stocks of DiNV. One stock of DiNV was isolated from infected cells and then passaged in Dinn-1 cells for 3 passages (DiNV Passage 4), and the other was DiNV semi-purified via sucrose density gradient ultracentrifugation and called “middle band”, which was a 5^th^ passage (MB DiNV) (Mulkey *et al.*, in prep). Dv-1 cells were counted with a hemocytometer and seeded into wells of two 48 well plates at a density of approximately 100,000 per well in 200 μL of growth medium. In each plate the interior 24 wells were seeded with the Dv-1 cells, and the outer wells were filled with DPBS to aid with evaporation control. The cells were allowed to settle and attach to the wells for 24 hours in a 23°C incubator before inoculation. The next day we made ten-fold serial dilutions of each virus fluid. Each plate would contain four wells that received no virus that serves as a cell control, and four wells received either 10^-1^, 10^-2^, 10^-3^, 10^-4^, or 10^-5^ dilution of the virus solution. Serial dilutions were made in Schneider’s *Drosophila* medium to an amount so that each well could receive 200 μL of solution. The seeded plates from the day before were dumped to remove the existing medium. Then, 200 μL of the appropriate dilution was added to each well containing Dv-1 cells. There was one 48 well plate for the P4 DiNV titration and the other plate was for the MB DiNV titration. Again, the surrounding outer wells in the plates were filled with DPBS to aid with evaporation control. The inoculated plates were incubated in a 23°C incubator for 48 hours.

After 48 hours the two cells were fixed with cold acetone and subsequently stained with the vp39 antibody. In brief, the medium from each plate was dumped off and replaced with ice cold 80% acetone (-20°C). The plates were incubated at -20°C for 30 minutes. Then, the acetone was dumped from each plate, and the plates were allowed to dry in open air for 20 minutes. After drying, the plates were stored at -20°C for another 24 hours until staining. Plates were taken from -20°C and allowed to come to room temperature. A 1% BSA, 0.1% Tween in 1X DPBS blocking buffer solution was gently added to each well and the plates were incubated with the buffer for 30 minutes at room temperature. After incubation, the solution was dumped out of the plates. A 1:1000 dilution of anti-vp39 rabbit antibody was made in 1% BSA, 0.1% Tween in 1X DPBS. 500 μL of the antibody solution was added to each well in the plates, and the plates were incubated in the dark for 1 hour. After incubation with the antibody, the plates were washed three times with DPBS. A 1:4000 dilution of goat anti-rabbit IgG Alexa Fluor 488 secondary antibody (Invitrogen #A-11008) was made in 1% BSA DPBS. 200 μL of secondary antibody solution was added to each well and incubated for 1 hour in the dark. After incubation with the secondary antibody, the plates were washed three times with DPBS. After the third wash, the plates were imaged with a Leica Thunder microscope with the 10X objective in bright field and 488 nm channels. Importantly, the same exposure and other settings were used for imaging the control and infected wells. For the MB plate, the fluorescent foci in in the 10^-3^ wells were counted for FFU titering. And for the P4 DiNV plate, the fluorescent foci in the 10^-5^ wells were counted by eye for FFU titering. The discrepancy here had to do with countability of the foci. The counts in each well were averaged to get an overall estimate of the number of FFU in the chosen dilution, and this number was used in the equation above to calculate the FFU/mL of the original stock solution.

### Determination of TCID_50_ with an endpoint dilution assay

We developed a basic end-point dilution assay to determine a 50% endpoint dilution titer expressed as the tissue culture infectivity dose that infects 50% of the inoculated cells (TCID_50_) per ml. The sample we titrated was DiNV P4 and we had quantitated this virus by qPCR to have a Cq value of about 14.5. We explored both ice cold 80% Acetone and 4% paraformaldehyde in DPBS as fixatives.

Dinn-1 cells were seeded into the interior wells of a 48 well plate in Schneiders *Drosophila* Medium (Gibco) containing FBS (Sigma, 10%) and gentamicin (Gibco, 10 μg/mL). Cells were incubated at 23°C in a non-humidified incubator until about 80% confluency (7 days). When the cells were at the desired confluency, using sterile technique, we dumped off the growth medium, blotted the plate on sterile paper towels and added 800 μL of Schneider’s Medium to each well. Next, we made 10-fold serial dilutions of our test virus in Schneider’s Medium with gentamicin by adding 0.5 mL of virus to 4.5 mL of diluent to make the 10^-1^ dilution. We repeated the dilution scheme to make 10^-2^, 10^-3^, 10^-4^, 10^-5^, 10^-6^ dilutions which were used for cell inoculation. We inoculated 3 plates of cells with 200 μL/well of appropriate dilutions and mock inoculated a cell control group.

Two plates were fixed with 80% acetone at 7- and 8-days post inoculation. The growth medium was removed by dumping and blotting with a paper towel. One mL/well of ice cold 80% acetone was added and incubated at -20°C for 20 minutes. The acetone was removed by dumping and blotting and the plate was air dried at room temperature for 30 minutes. 200μL of dilute (in DPBS) anti-p39 antibody was added to each well and incubated for 1 hour at room temperature. The plate was washed 3 times with DPBS and 100 μL per well of diluted goat ant-rabbit Alexa fluor 488 (Invitrogen) in DPBS was next added and incubated for 1 hour at room temperature in the dark. The conjugate was removed by dumping and blotting and 100 μL of water was added to each well. The plate was read using a Leica Thunder Fluorescent Microscope with a 488 nm filter.

The third plate was harvested 9 days post inoculation and fixed with 4% paraformaldehyde. The medium in the plate was dumped and the plate was blotted on paper towel. 500 μL of 4% PFA in DPBS was added to each well and incubated for 10 minutes at room temperature followed by 3 washes with DPBS. The cells were permeabilized with 0.05% Triton X100 in DPBS for 10 minutes at room temperature and washed 3 times in DPBS. The cells were then blocked with DPBS with 1% BSA, 22.52 mg/mL glycine, 0.1% Tween – 20. Then we added dilute anti-p39 in DPBS and incubated for 1 hour at room temperature followed by 3 washes with DPBS. Dilute goat anti-rabbit Alexa fluor 488 in DPBS was next added to the wells and incubated for 1 hour at room temperature followed by 3 washes in DPBS. Finally, water was added to each well. The plate was read using a Leica Thunder Fluorescent Microscope with a 488 nm filter.

The titer is expressed as TCID_50_/mL and is determined using the Spearman-Karber Method (42), log (TCID_50_) = log(highest dilution at which 100% infection is observed) – 0.5 + (total # positive wells / total # wells) + (The dilution factor for a 200 μL inoculum is + 0.7).

### Infection methods for adult flies

We established a protocol for DiNV systemic infections using a Nanoject II® from Drummond Scientific Company (PA, USA) (25, 43, 44). The other common mode of infection for *Drosophila* is a poke with a dissecting pin needle that has been dipped in an infectious solution (23, 45, 46). Both methods involve piercing the cuticle (usually in the thorax) of the fly while they are anesthetized on a carbon dioxide pad. A comprehensive infection protocol using nanoinjections was developed to standardize the method and to minimize the potential to cross-contaminate flies during the infection process. To protect against cross-contamination, a separate CO_2_ pad was designated for virus infection work only, and it was soaked in 10% bleach and rinsed and dried after every use. Additionally, all flies for infection purposes are handled exclusively on the bleach cleaned CO_2_ pad and maneuvered with autoclaved, single-use toothpicks. Flies were placed on the CO_2_ pad in a specific order; with control flies first, followed by infected flies, to limit transfer of virus through the pad surface. And if various dilutions of the virus were used, handling of flies who received higher dilutions (smaller dose) preceded those with lower dilutions (larger dose).

We pulled injection needles from the 3.5-inch capillary tubes supplied by Drummond (part 3-00-203-G/X) using a Sutter Instrument Co. P-30 needle puller set to 95 for Heat 1 and 78 for Heat 2. After the needle tip was broken, it was backfilled with mineral oil and then filled with the desired injection solution. The major incentive to use nanoinjector for virus infections is that it allows for a standardized delivery of fluid. A volume of 27.6 nL was tested on *D. innubila* and it did not appear to be a volume that was overwhelming to the size of the fly, and thus this volume is consistently used for all injection infections. Injections were performed using the toothpick to stabilize the back of the fly, and by piercing the fly in the thorax with the needle slightly below and anterior to the wing, then injecting the fluid.

DiNV infectious solution was grown in Dinn-1 cells and relatively quantitated with qPCR to a Cq value of 16. Single use aliquots were thawed for use and discarded afterwards, so that no experiment received a solution with more freeze-thaw cycles than another. To test the virulence of the 16 Cq DiNV cell lysate on nanojector injected *D. innubila*, two replicate blocks of infections were performed. *D. innubila* genome line (stock # TH190305) males aged 5-7 days post emergence were injected in their thorax with 27.6 nL of cell culture medium (10% fetal bovine serum supplemented Schneider’s Drosophila Medium, including penicillin, streptomycin, gentamicin, and amphotericin B) as control. For virus infection, male *D. innubila* aged 5-7 days post emergence were experimentally injected in their thorax with 27.6 nL of 16 Cq DiNV cell lysate. Flies were kept in standard mushroom food vials, with approximately 10 flies per vial, and transferred to fresh vials every 3 days. Flies that died within the first 24 hours or were lost during the experiment were removed from the analysis. Daily mortality was monitored for 12 days at which the experiment ended because all the experimentally infected flies had died.

For the infection experiment, the daily mortality data was analyzed in R Studio with the packages Survival and Survminer (47, 48). A custom R function was developed to convert the daily mortality data into the proper format for the analysis packages. The data was visualized via Kaplan-Meier plots and statistically summarized via a Cox proportional hazard model. Models comparing just infection or infection + block (replicate) were compared with Akaike Information Criterion (AIC) to determine whether block should be included in the model. The AICs were significantly different from each other, suggesting that the model with the smaller AIC, ∼infection + block, should be used as the appropriate model for the analysis.

### Comparing needle poke and nanoinjection

We aimed to test the magnitude and repeatability of dose differences using the nanoinjector compared to a needle poke. *D. innubila* males were either needle poked or injected with 16 Cq DiNV and frozen immediately to compare the dose of virus received by each fly. Three replicate flies were poked with a dissecting pin needle in the thorax that was dipped in 16 Cq DiNV between each fly. Three replicate flies were injected with 27.6 nL of 16 Cq DiNV in the thorax. Flies were individually frozen at -20°C within 1 hour of infection. DNA was extracted from each fly using the Qiagen Puregene kit in a modified protocol. In brief, individual flies in 1.5mL tubes were homogenized with a sterile pestle in 100 μL of chilled cell lysis solution on ice and then incubated at 65°C for 10 minutes. Samples were cooled to room temperature and 2 μg of RNase A solution was added to each tube. Samples were mixed by inverting 25 times and incubated for 40 minutes at 37°C. After incubation, 33 μL of protein precipitation solution was added to each sample, and samples were vortexed and incubated on ice for five minutes. Then samples were centrifuged for three minutes at 16,300 times g to pellet proteins and the exoskeleton. The supernatant was transferred to new 1.5 mL tubes containing 100 μL of fresh 100% isopropanol. Samples were inverted to mix 50 times and centrifuged at 16300 times g for five minutes to pellet the DNA. The DNA pellet was washed once with 70% ethanol, dried, and resuspended in 20 μL of DNA hydration solution. DNA was allowed to resuspend for a few hours at room temperature before being stored at -20°C while waiting for further processes.

To determine the difference in dose given to the flies when either needle poked or injected, the viral and host DNA was quantified with quantitative PCR (qPCR). DNA samples were diluted two ways for this analysis, either by diluting the DNA 1:10 or by diluting each sample individually to 1ng of DNA. Three technical replicates of each sample were run for qPCR with two primers. The *D. innubila* host gene primer *tpi* was used to quantify relative genome copy in the DNA sample (Dyer et al., 2005). The DiNV virus gene primer *PIF3* was used to quantify relative virus genome copy number in the DNA sample (see table S1 for primer sequences). Each reaction included 1 μL of DNA, 5μL of SsoAdvanced Universal SYBR Green Supermix (BioRad 1725270), 0.5 μL of 10 μM forward primer, 0.5 μL of 10 μM reverse primer, and 3 μL of molecular grade water. The qPCR program used included: An initial denaturation step of 3 minutes at 95°C, followed by 40 cycles of 10 second at 95°C and 30 seconds at 60°C, followed by a 5-minute 95°C extension. This is then followed by a melt curve section that increases by 0.05°C every 5 seconds from 55°C to 95°C.

Analysis of qPCR Cq results was performed in R. To calculate the relative amount of DiNV genome compared to fly genome as a single ΔCq calculation was first done by subtracting the average PIF3 Cq from the average tpi Cq for each sample. This was then raised as an exponent to 2 (2^ΔCq^). The 2^ΔCq^ is the relative amount of DiNV genome scaled to host genome DNA. First the mean of the 2^ΔCq^ for each infection type was compared to estimate the difference in dose given. Second, the coefficient of variation of the 2^ΔCq^ for each infection type was compared to estimate the difference in variation in dose given by the two methods.

### Testing for sex effects of DiNV infection

To test for differences in survival probability of male and female *D. innubila* at different DiNV dilutions, five different virus dilutions were made. DiNV grown in Dinn-1 cells (P4 DiNV) was quantified via qPCR to 14 Cq. This solution was titered with a Fluorescence Focus Unit (FFU) method and was about 2.38 x 10^2^ FFU/μL or 0.238 FFU/nL. For a 27.6 nL injection volume, this solution undiluted would deliver a total of 6.57 FFU to the fly. Based on this number, dilutions to deliver a total of 6, 3, 1, 0.1, and 0.01 FFU in 27.6 nL total volume were generated. Importantly, we do not think that 1 FFU equals 1 infectious unit, and that the FFU titer is a relative measure. Four replicate block injection experiments were performed to get enough replicates of the five P4 DiNV dilutions and the control cell culture medium for both male and female *D. innubila*. All flies were aged to 7-9 days post emergence to ensure that females were mated. Flies were kept in standard mushroom food vials, with approximately 10 flies per vial, and transferred to fresh vials every three days. Flies that died within the first 48 hours or were lost during the experiment were removed from the data. Daily mortality was monitored for 14 days at which time the experiment ended.

Daily mortality data was analyzed in R as described above. We compared the effect of sex, replicate block, and dilution as a continuous numeric variable. Three models were compared: ∼dilution + block + sex, ∼dilution + block, and ∼dilution + block + sex + dilution by sex interaction. AIC comparisons using the R package AICcmodavg suggested that the ∼dilution + block + sex model best fit our data which is what we used for analysis, however it only had slightly better predictive power than the other models. We also wanted to understand the effect of these variables on survival with only the four highest doses or the lowest dose and the control, so we used a reduced dataset of only the 0.1 FFU, 1 FFU, 3 FFU, and 6 FFU injected flies or only the 0.01 FFU and control injected flies. We again compared the same three linear models for each reduced dataset with AICcmodavg. For the 0.1 FFU, 1 FFU, 3 FFU, and 6 FFU injected flies a ∼dilution + block + sex model was the best fit and was used for analysis. For the 0.01 FFU and control injected flies the ∼dilution + block model was the best fit and was used for analysis.

### Validation of FFU DiNV titering method with nanoinjections

The fluorescence focus unit assay (FFU) was used to titer multiple different stocks of DiNV. To validate the titering method experimentally, we diluted a separate stock of DiNV to the same FFU doses that had been done with the P4 DiNV stock and injected them into male *D. innubila* and assayed survival and DiNV infection. The “middle band”, or MB, DiNV was titered to approximately 5.275 x 10^2^ FFU/μL and was diluted with cell culture medium to deliver either 0.01 FFU, 0.05 FFU, 1 FFU, or 3 FFU in a 27.5nL volume. Male *D. innubila* were infected in two replicate blocks, and all individuals were aged 5-7 days post emergence before infection and were given 27.6nL of solution. Flies were kept in standard mushroom food vials, with approximately 10 flies per vial, and transferred to fresh vials every three days. On transfer days, dead flies were frozen in individual 1.5mL tubes. Flies who died within the first 48 hours or were lost during the experiment were removed from the data. Daily mortality was monitored for 14 days at which the experiment ended, and all remaining alive flies were frozen individually. Infection status of all virus-treated flies from the male infection experiment (whether alive or dead) was determined by PCR amplification of *p47* as described previously (28) .

Daily mortality data was analyzed in R as described above. Cox proportional hazard models considering the dose of DiNV as a continuous variable were used to determine differences in survival. At first, we compared all doses with a ∼ dilution + block model, then we individually compared the 1 and 3 FFU doses, the 1 and 0.05 FFU doses, 0.01 and 0.05 FFU doses, and the 0.01 and CCM treatments separately for individual comparisons with a ∼ dilution + block model.

### Determining DiNV genome increase in various dilutions across early infection

A subset of FFU titer dilutions were selected to be used for injecting into *D. innubila* and measuring viral genome increase during the first few days of infection. Passage 4 DiNV was diluted with cell culture medium into 3 FFU, 0.1 FFU, and 0.01 FFU dilutions. Both male and female flies aged 5-7 days post emergence were injected with the various dilutions. Flies were frozen for DNA extraction on day 0 (within one hour of injection), day 1, day 3, and day 5 post injection. Only flies who were alive were frozen for analysis. In total, there were 8 flies for each sex, dilution, and day, a total of 192 flies. DNA was extracted from each fly individually as described above.

DNA was diluted 1:10 before use in qPCR reactions. Samples were not run with technical replicate qPCR reactions, instead employing 8 biological replicates. qPCR reactions were done for DiNV genome with the *PIF3* primers, and *D. innubila* host genome with the *tpi* primers.

Analysis of qPCR Cq results was performed in R. Relative viral genome increase was calculated with the ΔΔCq method. The first Δ was calculated by subtracting the average PIF 3 (viral DNA) Cq from the average tpi (host DNA) Cq for each sample, which compares DiNV genome to host genome. The second Δ was calculated by averaging the day 0 ΔCq for each dilution and sex and then subtracting that from the ΔCq for days 1, 3, and 5 for each dilution and sex. To obtain the fold increase, we raise 2 to the ΔΔCq power. In this way we calculated the relative genome increase from day 0 across the three timepoints. Relative viral genome increase (2^ΔΔCq^) and relative DiNV genome amount (2^ΔCq^) visualized with ggplot2 with box plots.

Statistical differences between relative genome increase were calculated with linear models in R. For all models the day and FFU dose were considered continuous variables. At first, we separated out the data between males and females. In each model we tested the effect on the non-transformed ΔΔCq and used the R package AICcmodavg to compare models. For each sex separately a model considering ∼day + dilution was analyzed. This model was a better fit than one including a day and dilution interaction. We additionally subset our data and just tested the effect of day on the 0.01 FFU dilution alone. We then combined the data from both sexes and compared multiple models with AIC. Surprisingly, this comparison suggested that the model that did not include sex as a variable, was the best fit for our data with the lowest AIC and that the other models do not add much more predictive power. Thus, we analyzed a ∼day + dilution + day by dilution interaction model with both sexes combined.

Infection status of all samples used in the qPCR experiment was determined by PCR amplification of a DiNV gene. DNA was extracted from each fly individually as described above. Each PCR reaction included 1μL of DNA, 5μL of GoTaq® Master Mix (Promega, M7123), 0.25μL of 10mM forward primer, 0.25μL of 10mM reverse primer, and 3.5μL of molecular grade water. The DiNV gene p47 was chosen to generate PCR primers for testing virus presence, and presence of DNA in each sample was validated with primers for host DNA, *CO1* and *tpi*. PCR products were run on a 1% agarose gel at 90V for 35 minutes and visualized under UV light. Samples were considered DiNV positive if they had a visible band at approximately 370bp for p47 and if samples had amplification for at least one of the control primers. DiNV infection status was analyzed in R by determining the proportion of individuals with the infection on each day and with each FFU dilution and visualized with bar plots in the ggplot2 package. Because there was no difference between males and females in the qPCR experiment, both males and females are combined for this analysis.

## 7. Results

### Two new cell lines developed

The process of a flask of cells transforming from primary cells to immortalized cells was different for the two species. A single flask of KG03447-derived cells showed remarkable cell division compared to all others. After about four months of the cells existing in one flask, where the medium had been refreshed multiple times, the cells were scraped and passaged to new flasks. For about 1 year these cells continued to grow and were regularly passaged when confluent. After that time, we considered the cells likely to be immortalized and the cells were consistently passaged at about a 1:10 split every two weeks using trypsin or cell scraping. To date they have been passaged over 40 times as well as frozen down, thawed and regrown. Following *Drosophila* Genomics Resource Center convention we named these cells E-Myd88[KG03447] and they are clearly a heterogeneous cell mixture (Fig 1A) - both round and elongated in shape. The cells can sometimes clump but mostly grow in a monolayer and can be easily counted with a hemocytometer.

**Figure 1.**
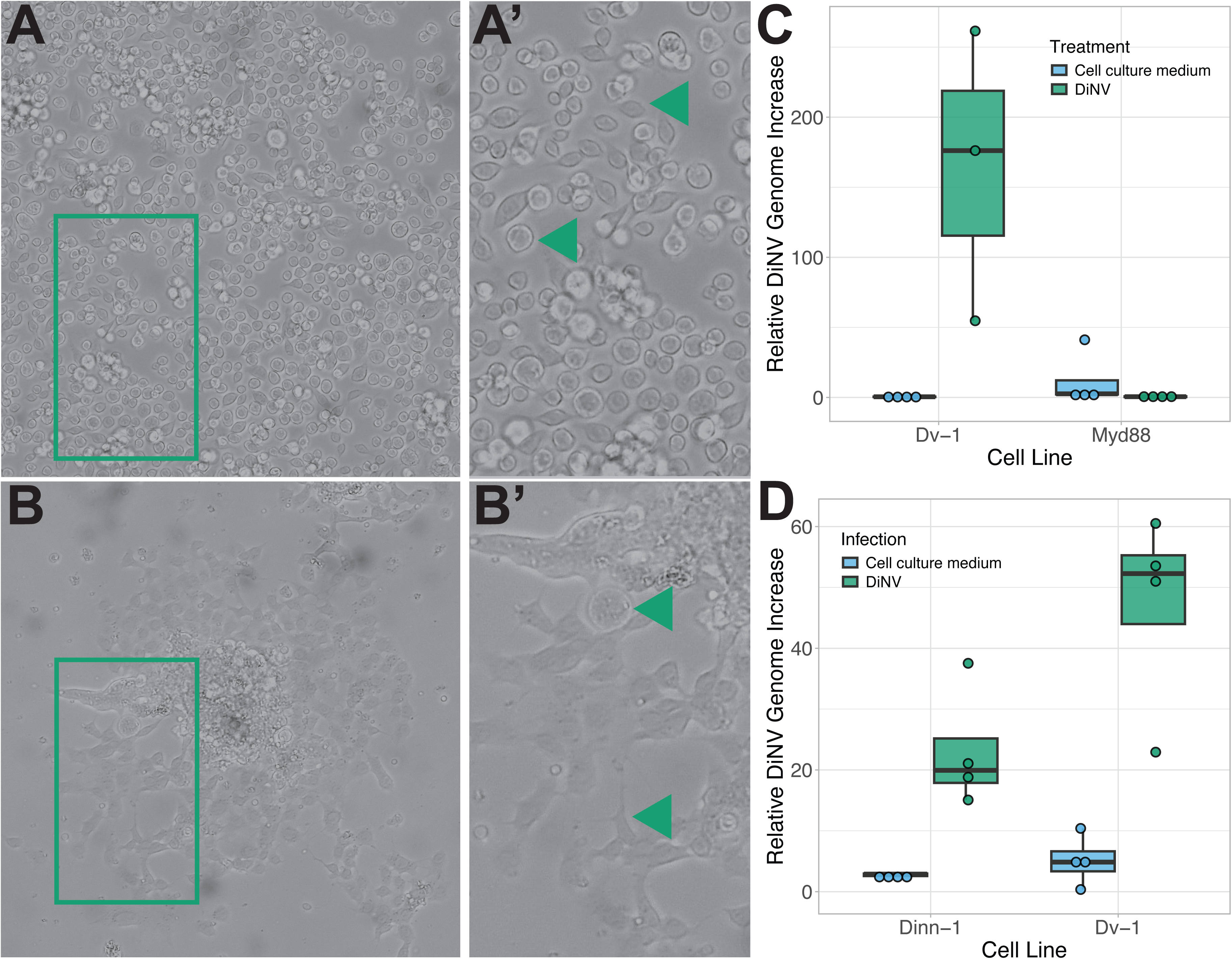
Representative images of E-Myd88[KG03447] and Dinn-1 cells. A) E-Myd88[KG03447] cells at passage 44. Box corresponds to the zoomed in image on the right, where the arrows indicate morphologically distinct cells. B) Dinn-1 cells at passage 17. Box corresponds to the zoomed in image on the right, where arrows indicate morphologically distinct cells. C) Relative DiNV genome increase from day 0 (2^ΔΔCq^) for Dv-1 or E-Myd88[KG03447] cells who were either given a control cell culture medium solution or inoculated with DiNV. Cells were incubated with the virus for five days. D) Relative DiNV genome increase from day 0 (2^ΔΔCq^) for Dv-1 or Dinn-1 cells who were either given a control cell culture medium solution or inoculated with DiNV. Cells were incubated with the virus for five days. ΔCq was calculated with absolute genome amount by comparison to Lambda DNA. In C and D, boxes represent the 25^th^ and 75^th^ percentiles, the solid line is the median and the whiskers show outliers (largest values no larger than 1.5 times the interquartile range).

The *D. innubila* cells took a much longer time to be considered immortalized and grow much slower than the E-Myd88[KG03447] cells. In fact, the flask of *D. innubila* primary cells that eventually immortalized was inoculated with a DiNV homogenate made from wild caught *D. innubila* flies. The primary *innubila c*ells inoculated with the DiNV containing fly homogenate did not die but continued to grow and a population eventually immortalized. This process took approximately 10 months and during this time, the original inoculated flask was repeatedly trypsinized and the cells were seeded to new flasks. All flasks seeded with the various passages of the Dinn cells except for one continued to produce DiNV positive fluids when assayed by qPCR. The DiNV negative Dinn cells were expanded, a frozen stock was made and labeled Dinn-1. These cells grow slowly, and they can be passed at a 1:3 split about every 2 weeks. Currently, approximately two years since the Dinn-1 cells were generated they have been passed over 20 times as well as frozen down, thawed and regrown.

Morphologically the Dinn-1 cells are very different from the E-Myd88[KG03447] cells (Fig 1B). The Dinn-1 cells are clumpy and grow out from large masses of cells where many cells are piled on top of each other. Surrounding these cell masses, a monolayer of cells grows out concentrically across the flask until it reaches the cells coming from another cell mass. These clumps are formed whether passing the cells with a cell scraper or with trypsin. The Dinn-1 cell line is a heterogeneous set of cells, where some are round and large, and others are very elongated and have multiple processes that spread out on the flask connecting cells together. The cells in the clump appear to mostly be round, while the cells forming a monolayer appear mostly to be elongated. Attempts to singularize these cells have so far been unsuccessful. For example, pushing the cells through a 26-gauge needle removes many of the larger clumps but still retains small ones. Additionally, the cells after the needle shearing do not grow very well.

The newly developed cell lines can be transfected with a lipofectamine based method and are able to express GFP under a promoter from *D. melanogaster* (Fig S1). E-Myd88[KG03447] cells (Fig S1 right) show the most expression of GFP after 5 days post transfection, where many cells show strong green signals. For this experiment we also used cells derived from *Drosophila virilis* larvae (WR Dv-1). Dv-1 cells (Fig S1 left) are sparsely transfected, as are Dinn-1 cells (Fig S1 middle), yet they are clearly transfected. GFP positive E-Myd88[KG03447] cells were visible after 24 hours post transfection. GFP positive Dv-1 and Dinn-1 cells were visible after 48 hours, although very few, and more became visible after subsequent days. While the transfection efficiency of the Dinn-1 cells appears to be quite low, this experiment shows that not only can the Dinn-1 cells be transfected with a broadly available reagent, but that in future experiments promoters from *D. melanogaster* may be used.

### Dinn-1 cells but not Myd88 cells allow for viral genome replication

*D. innubila* (Dinn-1) cells replicate the DiNV genome and *D. melanogaster MyD88* mutant (E-Myd88[KG03447]) cells do not (Fig 1C-D). The closer relative *D. virilis* (Dv-1) cells also replicate the DiNV genome. Figure 1C shows the relative DiNV genome increase after 5 days for Dv-1 and E-Myd88[KG03447] cells infected with DiNV. The DiNV primer was normalized to Lambda DNA as a DNA extraction control to account for the different numbers present in samples from different cell lines. This gives us an absolute increase in genome amount. The DiNV genome increased by 164-fold in Dv-1 cells. However, for E-Myd88[KG03447] cells infected with DiNV we see no DiNV genome increase, indicating that these cells are not able to replicate the virus genome. With a linear model we see that infection status increases DiNV growth (p = 0.038), but that there is a significant interaction effect between cell type and infection status (p < 0.00001) because the E-Myd88[KG03447] cells do not grow DiNV. In contrast, the Dinn-1 cells do replicate the DiNV genome (Fig 1D). For infected Dv-1 cells on average the DiNV genome increased by 12-fold. For infected Dinn-1 cells on average the DiNV genome increased by 9-fold. With a linear model we see that infection status increases DiNV growth (p = 0.00003), however for this data the best model was one that did not include cell type as a variable, indicating that DiNV growth is not different between the Dinn-1 and Dv-1 cells.

If we use host cell primers to normalize the viral replication for the Dv-1 and Dinn-1 cells, this gives us an increase in viral genome amount per host genome, which we used to account for the likely different number of cells between the two cell types. Using this method, we see slightly different results (Fig S2), where infection status also increases DiNV growth (p = <0.00001), cell type influences DiNV growth (p = 0.0395), and that there is a significant interaction between cell type and infection status (p = 0.0105). We do know that we cannot standardize the number of cells between Dv-1 and Dinn-1 cells in these experiments because we cannot count Dinn-1 cells, thus we attempt to control for this by also comparing viral genome number to host genome.

### Anti-DiNV vp39 antibody specifically binds to a 29 kilodalton band in a western blot of DiNV Passage 5 fluids

A western blot of Dinn-1 cell and DiNV Passage 5 fluids separated on a 10% SDS-PAGE gel was stained with mouse monoclonal anti-actin and rabbit polyclonal anti-vp39 antibodies produced a positive band at about 43 kD (actin) in both samples and a band at about 29 kD (vp49 nucleocapsid) in only the DiNV containing samples (Figure S3).

### A fluorescence focus-forming unit assay (FFU) for quantifying infectious viral particles

The FFU assay was preliminarily tested on Dinn-1 and Dv-1 cells using fluorescent antibody detection of the DiNV vp39. First, we discovered that the clumpiness of Dinn-1 cells causes some issues with visualizing fluorescence at the low magnifications needed for the assay. The clumps auto-fluoresce in a way that can make it hard to see foci of infection (Fig S4). Previous work using indirect fluorescence antibody staining and confocal microscopy on DiNV infected Dinn-1 and Dv-1 cells using the same DiNV anti-nucleocapsid antibody showed us that DiNV is sequestered in cells to multiple puncta inside the cell (Mulkey *et al.* in prep). At the low magnification (10X) used for visualizing the FFU assay, the puncta are seen as very small dots of fluorescence (see inset in Fig S4). The elongated shapes of some of the Dinn-1 cells make it hard to know where one cell begins and another ends, thus making it difficult to know if a punctum is in one cell or another without staining for cell membranes or other cellular structures. However, in Dv-1 cells, we can determine whether puncta are in one cell or two and are somewhat easy to visualize (Fig S4B). We were able to titer the DiNV stock (passage 4 DiNV) to approximate 2.37 x 10^5^ FFU/mL with the FFU method on Dv-1 cells. We note, however, that Dv-1 cells from *D. virilis* are quite divergent from the native *D. innubila* host and so the number of focus-forming units in Dv-1 cells may not accurately represent the number of infectious viral particles in *D. innubila* (flies or cells). We treat FFU as a relative measure of infectious viral particles.

### An end-point dilution assay to determine viral titer

We titrated a passage 4 sample of adapted DiNV to determine TCID_50_/mL, the dose required to infect 50% of tissue culture samples. We harvested one plate at day 7, day 8 and day 9 post inoculation. The day 7 and day 8 plates were fixed with 80% acetone, and the day 9 plate was fixed with paraformaldehyde. The TCID_50_/mL of the two plates fixed with acetone were 10^4.95^ (∼8.91 x 10^4^) infectious units/mL (day 7) and 10^5.2^ (∼1.58 x 10^5^) infectious units/mL (day 8, Table S2). The titer of the day 9 plate fixed with paraformaldehyde was 10^5.95^ (∼8.91 x 10^5^) infectious units/mL. Thus, the results of the fluorescence focus-forming unit (2.37 x 10^5^ FFU/ml) and end point dilution (8.91 x 10^4^ to 8.91 x 10^5^) assays are comparable.

### Infection of whole flies leads to rapid mortality

First, we aimed to determine whether injection with DiNV caused noticeable mortality in male *Drosophila innubila*. Indeed, *D. innubila* injected with DiNV passage 3 with a qPCR Cq value of 16 die much faster than flies injected with cell culture medium (Fig 2A). By 12 days post injection, all flies that received the DiNV treatment died, while only one control fly died (Cox PH p = 1.55 x 10^-11^). The rapid and complete mortality of DiNV infected flies is different than previous infections with *D. innubila* that used a dissecting needle poke for virus delivery where the maximum mortality after 15 days was ∼75%, however the virus inoculum used was different and titer was less controlled, thus the results cannot be compared directly (Hill and Unckless, 2020). However, this does suggest that a simple needle dipped in virus solution and poked may not be sufficient to deliver enough virus to the host to cause complete and rapid death. The utility of the injection method compared to the poke method was investigated further by quantifying the amount of virus given to a fly at the time of infection.

**Figure 2.**
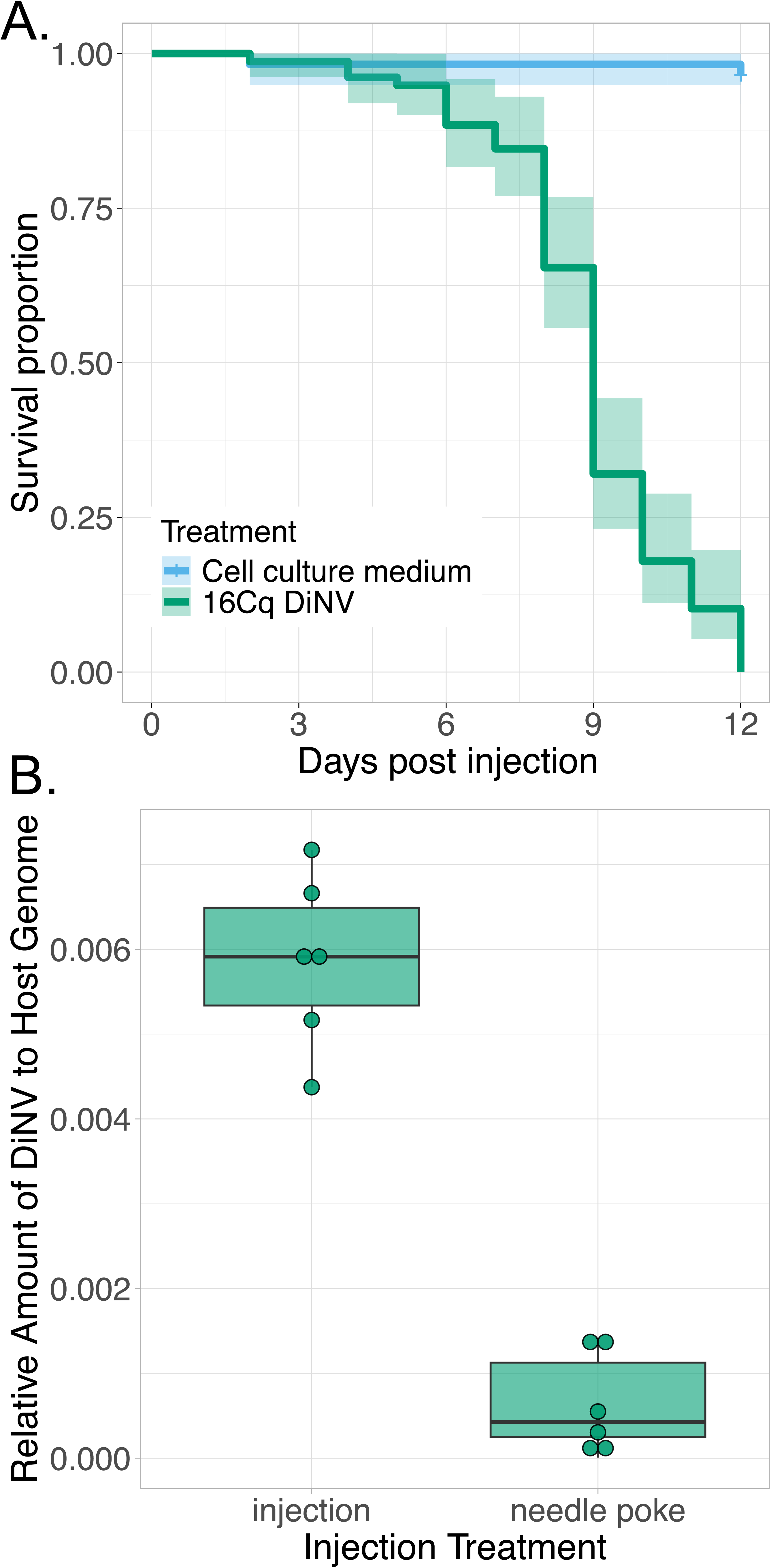
Preliminary nanoinjection investigations. (A) Kaplan-Meier survival curve for infection with DiNV. Male *D. innubila* were either injected with cell culture medium or 16Cq DiNV solution and survival was monitored for 12 days. Lines represent the average survival proportion across all replicates. (B) Dotplot comparing relative amount of DiNV DNA to host DNA (ΔCq) between *D. innubila* flies injected with 27.6nL of DiNV or flies poked with a needle dipped in DiNV. Dots represent an individual fly. Error bars represent standard error of the mean. The DNA was diluted in two ways as input into qPCR, either to 1ng or 1:10. In B, boxes represent the 25^th^ and 75^th^ percentiles, the solid line is the median and the whiskers show outliers (largest values no larger than 1.5 times the interquartile range).

Comparison of qPCR of DNA of *D. innubila* infected with DiNV either with a needle poke or a 27.6nL injection reveals that the injection delivers far more virus to the flies (Fig 2B). The average relative amount of virus compared to host genomes (2^ΔCq^) for the needle poke was 0.00055, while the average for the injection was 0.0057. This indicates that the injection delivers approximately 10x the amount of virus than the needle poke. Perhaps more importantly, the coefficient of variation (CV) for viral titer directly after needle poke was 1.374 compared to 0.077 for injections, indicating that the amount of virus DNA given with the needle poke is approximately 18 times more variable than with injection.

### No evidence for differential mortality in males and females

Surprisingly, most dilutions of DiNV tested resulted in complete mortality of all flies by day 14 in both sexes (Fig 3). Only the 0.01 FFU in both sexes, and the 0.1 FFU in females resulted in intermediate mortality. There was also very little variation in the slope of the survival curves for the dilutions that resulted in complete mortality. For males, the median survival time was 8 days for 6 FFU and 3 FFU, 8.5 days for 0.1 FFU, and 9 days for 1 FFU (Fig 3A). For females, the median survival time was 8 days for 6 FFU, 1 FFU, and 0.1 FFU, and was 7 days for 3 FFU (Fig 3B). When considering all treatments and both sexes, the dose of DiNV (p = 8.2×10^-16^), replicate block (p = 0.0247), and sex (p = 0.0412) had a significant effect on survival proportion after infection. Though the sex effect appears small, males have a more gradual survival curve for some of the dilutions. To further tease out these effects we compared linear models with subsets of dilutions.

**Figure 3.**
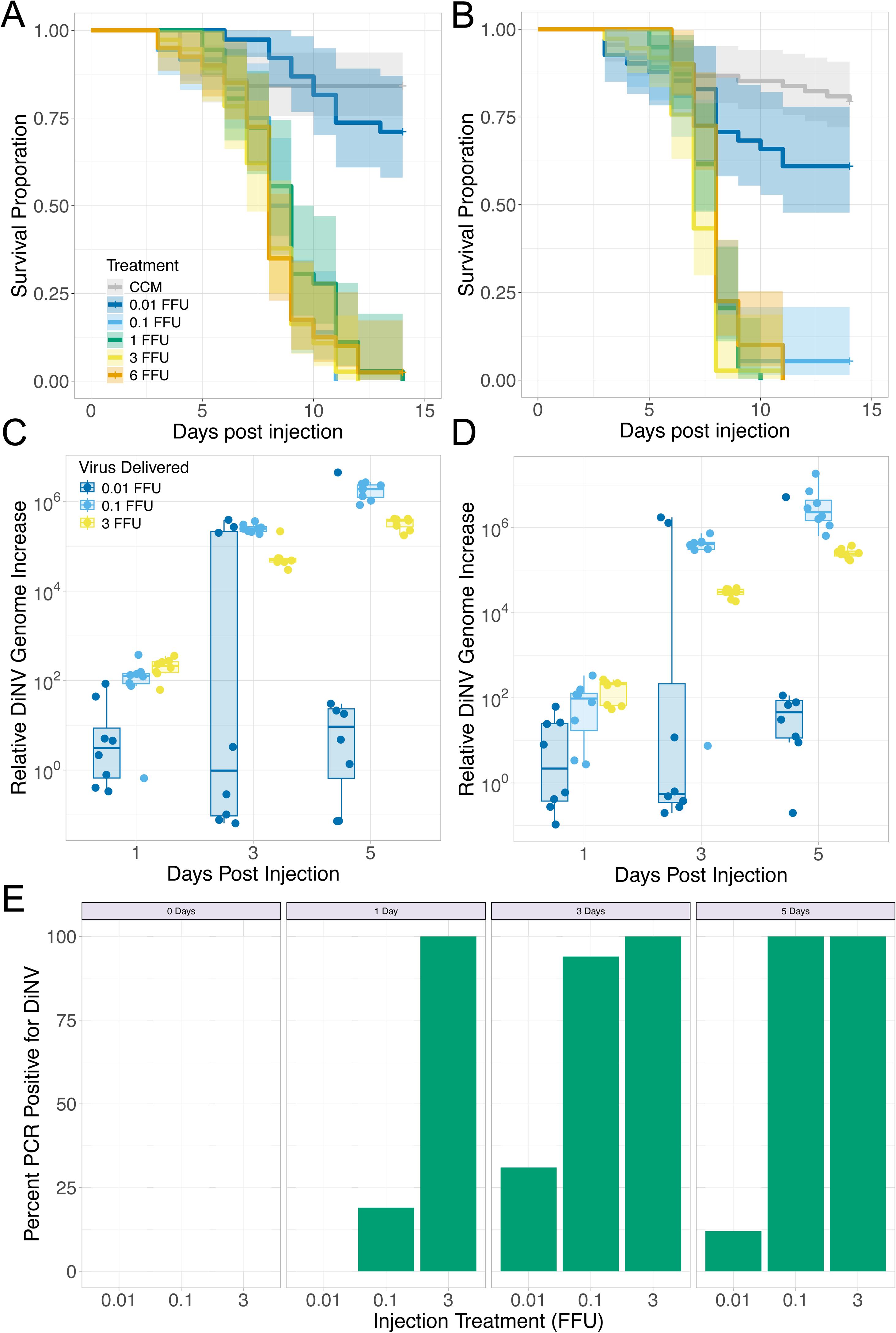
Comparing male and female *D. innubila* during DiNV infection. Kaplan-Meier survival curve for infection with different dilutions of DiNV. Male (A) and female (B) *D. innubila* were either injected with cell culture medium, 6 FFU, 3 FFU, 1 FFU, 0.1 FFU or 0.01 FFU of P4 DiNV and survival was monitored for 14 days. Lines represent the average survival proportion across all replicates. Box plots showing relative DiNV genome increase from day 0 (ΔΔCq) for females (C) and males (D) injected with either 3 FFU, 0.1 FFU, or 0.01 FFU P4 DiNV. Boxes represent the 25^th^ and 75^th^ percentiles, the solid line is the median and the whiskers show outliers (largest values no larger than 1.5 times the interquartile range). E) Bar plots showing the percent of individuals positive for a DiNV PCR marker from the experiment in A and B, males and females for each treatment and day are combined. Numbers inside each bar correspond to the number of individuals with each PCR result (yes or no).

When we only consider the four lowest dilutions, (6 FFU, 3 FFU, 1 FFU, and 0.1 FFU), dilution does not have a significant effect on survival (p = 0.769), but sex does (p = 0.00002). This is likely because females have a slightly more rapid decline in survival for these doses than the males. What this suggests is that for the highest doses there is no difference in survival even if we dilute the virus over tenfold (6 FFU compared to 0.1 FFU) when we are starting with such high doses. In contrast, when considering only the 0.01 FFU dilution and the control, the injection treatment does have a significant effect on survival (p = 0.0185), but sex is not significant. Thus, sex appears to have much less influence on survival for DiNV than it does for the related Kallithea virus in *D. melanogaster* (25).

### Validation of FFU DiNV titering method with nanoinjections

We validated our fluorescence focus unit (FFU) assay by titrating two different stocks of DiNV and diluting them to the same FFU concentrations and then monitoring mortality in flies infected with the two DiNV stocks. Middle band (MB) DiNV was titrated with the FFU method and diluted to deliver 0.01 FFU, 0.05 FFU, 1 FFU, and 3 FFU and injected into male *D. innubila*. In general, the survival curves of MB DiNV infected flies (Fig S5) are like the survival curves of P4 DiNV infected flies at the same dilutions (Fig 3). As expected, dose of DiNV has an overall significant effect on mortality (p = <0.00001), and replicate block does not (p = 0.355). There is no significant difference in mortality between the 1 and 3 FFU MB DiNV treatments (p = 0.089), which is the same as what was seen with P4 DiNV. However, there is a significant difference between the 0.05 FFU and 0.01 FFU MB doses (p = <0.00001), as well as the 0.01 FFU MB and CCM treatments (p = 0.0121). These different doses also cause significant effects on survival for diluted passage level 4 DiNV.

General trends with this infection experiment also mirror the previous experiments. The median survival time for males infected with MB DiNV at 0.05 FFU was 9 days, which is the same as males infected with P4 DiNV. Median survival time for MB DiNV at 1 FFU and 3 FFU was 9 and 8 days, respectively. This is very similar to the median survival time for P4 DiNV at these doses; 8 and 7 days. Importantly, no individual is alive past day 12 with either dose or with either DiNV stock. Although the survival curves cannot be compared directly because the experiments were done at different times, the curves are very similar across DiNV stocks and indicate that the virus was likely diluted to a similar amount in each experiment.

To further test the method, PCR on each infected fly was performed to assess infection status. All individuals who received a 1 or 3 FFU dose of MB DiNV were infected with the virus. Only one individual who received the 0.05 FFU MB DiNV dose did not show infection when alive at day 14, which is the same as what we saw for the P4 DiNV dilutions (Fig S5). And a little over half (26) of individuals given 0.01 FFU of MB DiNV did not show virus infection. This is like the P4 DiNV 0.01 FFU dose where a little over a third of individuals did not show virus infection. These results, along with the similarities in survival curves between the different viral stocks, suggest that the FFU titration method provides a reproducible and standardized way to quantify the amount of DiNV.

### DiNV genome increases in various dilutions across early infection

The DiNV genome copy number (a proxy for DiNV replication) increases greatly in the first 5 days post infection for both males and females (Fig 3C-D). When considering the sexes separately in linear models we see the same effects on DiNV growth. For males and females, as the days post infection increase and the dose increases, the relative DiNV growth increases. Both day post infection and dose cause significant change in relative increase in DiNV for males (p = 0.00002 and p = 0.00741 respectively), as well as females (p <0.00001 and p = 0.0134 respectively). By one day post injection, the 0.1 and 3 FFU flies have an average increase in DiNV, which further increases until day five. On day 3 and day 5 in both sexes, on average flies given the 0.1 FFU dose show a greater increase in DiNV than the flies given the 3 FFU dose (Fig 3C-D). However, when we look at relative amount of DiNV genome compared to host genome (Fig S6), these two dilutions have roughly the same titer. The difference in genome increase is likely caused by the different initial doses of DiNV genome given; to get to the same relative amount of DiNV as the 3 FFU treatment on day three or day five, the 0.1 FFU flies must grow more virus.

When we consider a linear model with males and females combined, our analysis indicated that a model using sex as a variable overparameterized our data. This tells us that there is no difference in DiNV genome increase between the males and females in any of our variables.

Flies given the 0.01 FFU dose show a different pattern than the flies given the 0.1 FFU and 3 FFU doses (Fig 3C-D). When we subset out just the individuals who received this low dose, there is no significant increase in viral growth over the five-day experiment for either males or females (p = 0.604 and p = 0.18 respectively). Some individuals do not show any growth of the virus, while some other individuals have viral increase that is comparable to the higher doses, which is why we cannot detect any statistical effect of day for this dilution. This led us to check each sample with standard PCR for viral presence at each time point (Fig 3E). We see that most of the individuals with the 0.01 FFU dose do not show any presence of DiNV infection through day five, while all individuals are infected by day five with the higher doses. We also see that using PCR is an imperfect way to detect whether an animal is infected initially, because we know that all individuals with the 0.1 or 3 FFU doses will end up dying from the virus but come up as virus negative after injection (day 0). However, with the inability to sample DNA from a fly over time, this is still one of our best methods to estimate whether a fly is infected or not. It is not clear whether some individuals given the 0.01 FFU dose do not actually receive infectious virions, and thus never have any viral growth, or if the five-day timepoint is too early to detect if all samples do end up growing virus.

## 8. Discussion

The development of two new cell lines, and the *D. innubila* cells in particular, is a major advancement for the DiNV host-virus model system. The Dinn-1 cells allow us to grow DiNV in the lab instead of relying on wild caught infected flies as a virus source. Notably, all previous published research on DiNV has been based on a wild caught source of the virus, which can be limiting because field collections are not always feasible. While we do see some preliminary evidence of DiNV adaptation to cells in culture, we do not see evidence of virulence attenuation. DiNV passaged in Dinn-1 cells is very virulent and can cause complete mortality in experimentally injected flies even at a high dilution. Thus, the Dinn-1 cells have already been instrumental in aiding DiNV research.

Dinn-1 cells can be used in experiments on DiNV infection that are less tractable in whole animals. It is much easier to deliver chemical inhibitors to cells than to whole flies (49), and so experiments that aim to block host cellular processes, such as apoptosis, are now possible with Dinn-1 cells. Additionally, specific host genes can be inhibited by delivery of small interfering RNAs (RNAi) in cell culture (50), and this is especially important with this system because we do not have the ability to drive the expression of hairpin RNAs in *D. innubila* like what is possible in *D. melanogaster* with the GAL4-UAS system (51). Future experiments could knock down expression of host immune genes and look for a change in DiNV genome replication or increase in viral titer to see what immune pathway has the most effect on DiNV. The infection dynamics and gene expression profiles of infected cells versus whole animals, or infection across time can be investigated with RNAseq. The list of potential experiments now possible with Dinn-1 cells is vast and ever growing, and importantly the development of the Dinn-1 cell line has opened many areas of possible research for the DiNV host-virus system.

The nanoinjector technology for systemic DiNV infections opens many doors to many questions about the dynamics of DiNV infection, not only in *D. innubila*, but potentially in multiple species of *Drosophila*. Many species of *Drosophila* in the wild, from *D. tenebrosa* to *D. melanogaster*, have been associated with DiNV (28). However, very little has been done to experimentally infect multiple species of *Drosophila* with DiNV in the laboratory under more controlled conditions. One small experiment looked at *D. falleni*’s response to infection with DiNV in terms of mortality (28), however there was no investigation into viral load in this species, or if different viral titers affect mortality. A systematic investigation of the infection dynamics of DiNV across species of *Drosophila* is now possible. The modulation of injection volume used on differently sized flies is possible, and one can adjust viral titer dilutions to deliver consistent doses with differing volumes. Additionally, an even more preliminary experiment should be investigated. All experimental infections above were done with the *D. innubila* genome line, and while there is little population structure in wild *D. innubila* (52) different isolated mountain ranges in the Arizona Sky Islands harbor populations of *D. innubila* infected with DiNV (29). Different lines of *D. innubila* from different Sky Islands should be challenged with DiNV and their infection susceptibility investigated.

With the development of a systemic infection model, many other questions can be investigated. The tissue tropism of a systemic infection can be elucidated by injecting flies with DiNV, and then fixing, sectioning, and staining them with a DiNV specific antibody (anti-vp39) to understand what tissues DiNV infects. It is thought that DiNV is transmitted fecal-orally in the wild, however with a systemic infection the virus is transmitted throughout the body quickly via the hemolymph. So, even with a systemic infection we should be able to see what tissues DiNV is able to infect. To understand the host response to DiNV infection, an investigation into the transcriptional response to the virus with RNASeq should be undertaken. Previous research on DiNV has looked at RNA differential expression from wild caught male *D. innubila*, however the dose and time since infection with DiNV is unknown in these samples (29). The ability to deliver a consistent dose of DiNV allows for rigorous infections that can be investigated with multiple transcriptomic methods, be it bulk RNASeq, small RNASeq, or even single cell RNASeq. The development of this nanoinjection method has opened many doors for expanding DiNV and *D. innubila* into an established model system for host-pathogen interaction.

## 10 Author statements

### 10.1 Author contributions

Conceptualization: MES, KMM, RLU; Data curation: MES, RLU; Formal analysis: MES, KMM, RLU; Funding acquisition: RLU; Investigation: MES, KMM; Methodology: MES, KMM, RLU; Project administration: RLU; Visualization: MES, RLU; Writing: MES, RLU,KMM

### 10.2 Conflicts of interest

The authors declare that there are no conflicts of interest.

### 10.3 Funding information

Funding for this project comes from NSF EAGER 2135167 to RLU and a University of Kansas Office of Graduate Studies Summer Research Scholarship to KMM.

## 10.4 Acknowledgements

We thank members of the Unckless lab – particularly Taiye Adewumi and Kistie Brunsell for helpful advice and feedback, Kelly Dyer, Paul Ginsberg, Brandon Cooper and Tom Hill for fly collections. Brian Ackley helped with microscopy. We acknowledge *Drosophila* Genomics Resource Center (NIH Grant 2P40OD010949) for providing cell lines. Funding was provided by NSF Grant 2135167 to RLU and a University of Kansas Office of Graduate Studies Summer Research Scholarship to KMM.

## Notes

### Competing Interest Statement

The authors have declared no competing interest.

