## Supplemental Materials for "The study of the *Drosophila innubila* Nudivirus in cells and flies"

**Supporting Tables**

| **Primer name** | **Sequence** |
| --- | --- |
| p47 F | TGAAACCAGAATGACATATATAACGC |
| p47 R | TCGGTTTCTCAATTAACTTGATAGC |
| CO1 F | GGTCAACAAATCATAAAGATATTGG |
| CO1 R | TAAACTTCAGGGTGACCAAAAAATCA |
| Lef4 F | CTTGGGGGCTACTTTCGACA |
| Lef4 R | TGGAAGAATGTTCCAATGGGGT |
| Lef9 F | CCAAGAATTTGGCGCGAACA |
| Lef9 R | GTTTGCGTCCATTGCGACTT |
| tpi F | GCCGGACAGAATGCCTACAA |
| tpi R | CCCAATCGGCGCCAAT |
| 16S F | AGAGTTTGATCCTGGCTCAG |
| 16S R | CGGTTACCTTGTTACGACTT |
| RPL11 F | GCAGCCCGTGTTTTCTAAGG |
| RPL11 R | TACTCGCGAACTTTCAAGCCAC |
| Lambda F | CGGCGTCAAAAAGAACTTCC |
| Lambda R | GCATCCTGAATGCAGCCATA |
| PIF3 F | GGATCGCCAAGCAAATAGGC |
| PIF3 R | GCCGTGAACCTGGTGTTTTT |

**Table S1.** Table of PCR primer names and sequences

|  | **dpi** | **10^-3^** | **10^-4^** | **10^-5^** | **10^-6^** | **TCID_50_/mL** |
| --- | --- | --- | --- | --- | --- | --- |
| Plate 1 - Acetone | 7 | 4/4 | 2/4 | 1/4 | 0/4 | 10^4.95^ |
| Plate 2 - Acetone | 8 | 4/4 | 3/4 | 1/4 | 0/4 | 10^5.2^ |
| Plate 3 - PFA | 9 | 4/4 | 4/4 | 2/4 | 1/4 | 10^5.95^ |

**Table S2. End-point dilution assay results.** Ratio of positive wells per dilution (top row) and calculated TCID_50_ per mL of inoculum for each plate. dpi: days post inoculation.

**Supporting Figures**

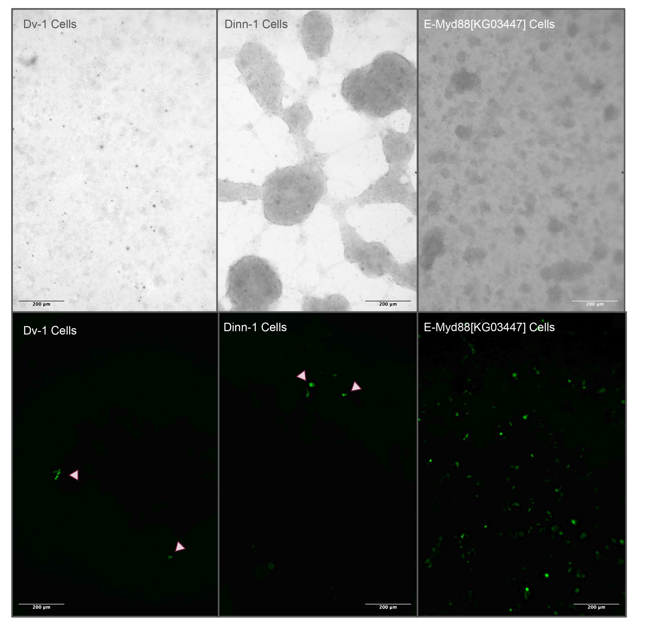

**Figure S1. GFP expression plasmid transfection of cell lines**. Cells were transfected with pAc5.1B-EGFP plasmid DNA and imaged five days later. The top panel is a bright field view of the cells while the bottom panel is GFP fluorescence. Arrows point to GFP expressing cells in the Dv-1 and Dinn-1 panels.

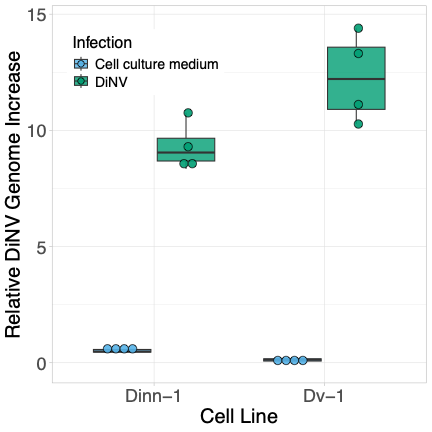

**Figure S2: DiNV infection in Dv-1 and Dinn-1 cells.** Relative DiNV genome increase from day 0 (2^ΔΔCq^) for Dv-1 or Dinn-1 cells who were either given a control cell culture medium solution or inoculated with DiNV. Cells were incubated with the virus for five days. ΔCq was calculated by comparing DiNV DNA to host DNA.

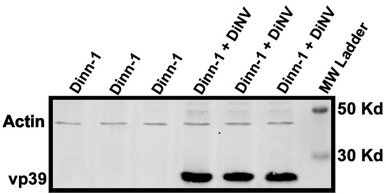

**Figure S3: Western blot of anti-DiNV vp39 antibody and Actin.** Three biological replicates each of Dinn-1 cells either mock-inoculated or inoculated with DiNV, then harvested and stained 10 days after inoculation.

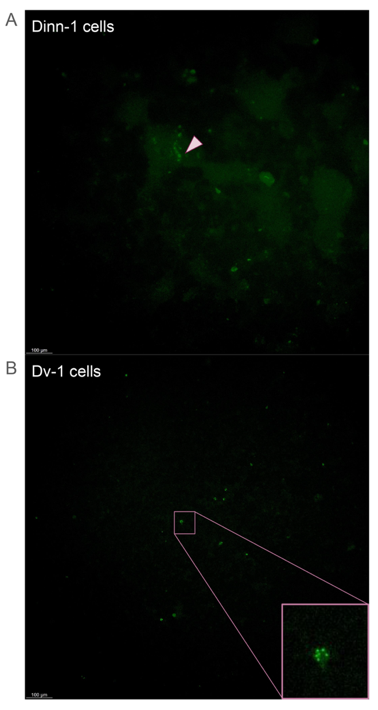

**Figure S4. Preliminary FFU titering of Dinn-1 and Dv-1 cells.** Dv-1 and Dinn-1 cells were inoculated with serially diluted P4 DiNV. Then the cells were fixed and stained with the DiNV P39 antibody. (A) The Dinn-1 cells were incubated for 24 hours before staining and a 10^-2^ dilution is pictured. Arrows point to DiNV positive cells. (B) The Dv-1 cells were incubated for 48 hours before staining and a 10^-3^ dilution is pictured. Inset is a DiNV positive cell with puncta visible. Images are GFP fluorescence.

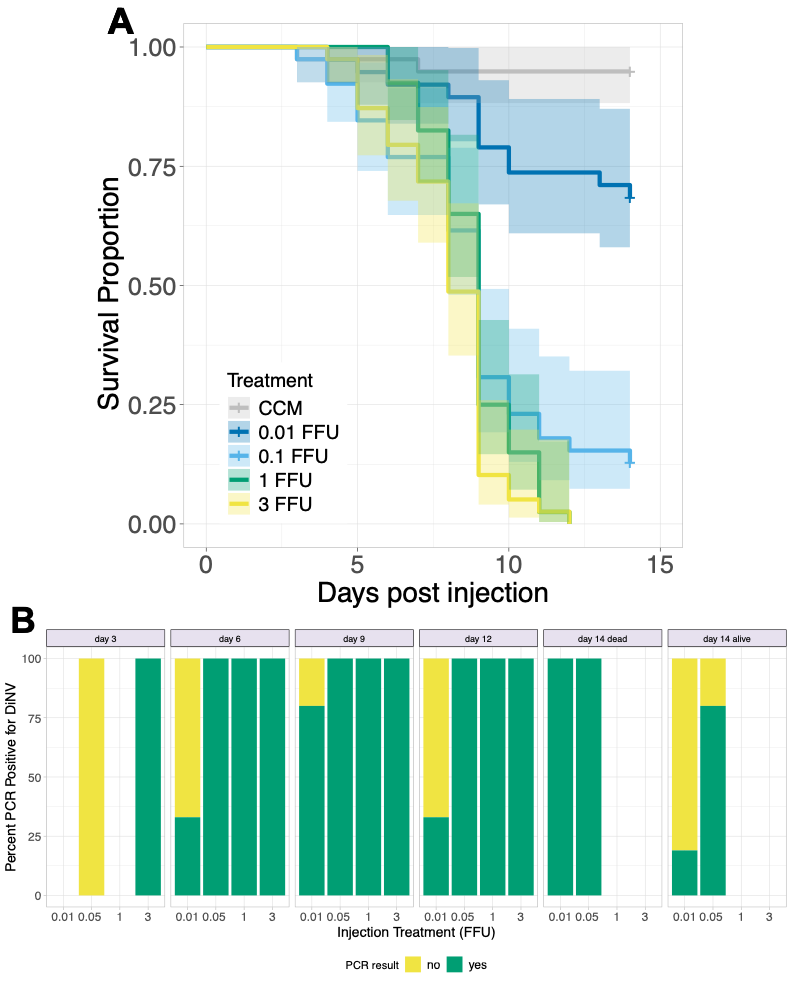

**Figure S5. Dilution infections of a different stock of DiNV in male *D. innubila*.** (A) Kaplan-Meier survival curve for infection with different dilutions of DiNV. Male *D. innubila* were either injected with cell culture medium, 3 FFU, 1 FFU, 0.05 FFU, or 0.01 FFU of MB DiNV and survival was monitored for 14 days. Lines represent the average survival proportion across all replicates. (B) Bar plots showing the percent of individuals positive for a DiNV PCR marker from the experiment in A. Numbers inside each bar correspond to the number of individuals with each PCR result (yes or no).

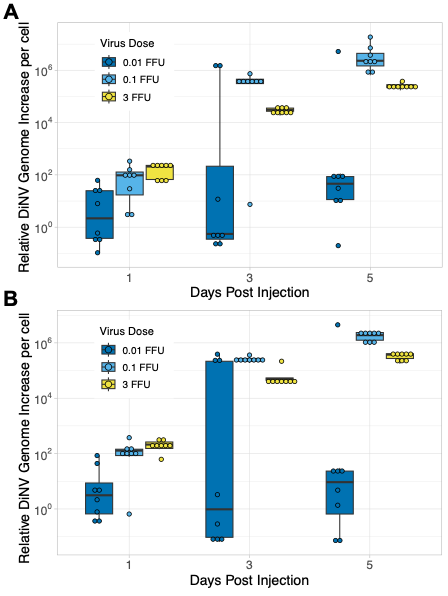

**Figure S6. Relative amount of DiNV DNA in hosts during early infection.** (A-B) plots showing relative DiNV genome (2^ΔCq^) for females (A) and males (B) injected with either 3 FFU, 0.1 FFU, or 0.01 FFU Passage 4 DiNV.
